# Motor-targeted spinal stimulation promotes concurrent rebalancing of pathologic nociceptive transmission in chronic spinal cord injury

**DOI:** 10.1101/2023.04.12.536477

**Authors:** Maria F. Bandres, Jefferson L. Gomes, Jacob G. McPherson

## Abstract

Electrical stimulation of spinal networks below a spinal cord injury (SCI) is a promising approach to restore functions compromised by inadequate excitatory neural drive. The most translationally successful examples are paradigms intended to increase neural transmission in weakened yet spared motor pathways and spinal motor networks rendered dormant after being severed from their inputs by lesion. Less well understood is whether spinal stimulation is also capable of reducing neural transmission in pathways made pathologically overactive by SCI. Debilitating spasms, spasticity, and neuropathic pain are all common manifestations of hyperexcitable spinal responses to sensory feedback. But whereas spasms and spasticity can often be managed pharmacologically, SCI-related neuropathic pain is notoriously medically refractory. Interestingly, however, spinal stimulation is a clinically available option for ameliorating neuropathic pain arising from etiologies other than SCI, and it has traditionally been assumed to modulate sensorimotor networks overlapping with those engaged by spinal stimulation for motor rehabilitation. Thus, we reasoned that spinal stimulation intended to increase transmission in motor pathways may simultaneously reduce transmission in spinal pain pathways. Using a well-validated pre-clinical model of SCI that results in severe bilateral motor impairments and SCI-related neuropathic pain, we show that the responsiveness of neurons integral to the development and persistence of the neuropathic pain state can be enduringly reduced by motor-targeted spinal stimulation while preserving spinal responses to non-pain-related sensory feedback. These results suggest that spinal stimulation paradigms could be intentionally designed to afford multi-modal therapeutic benefits, directly addressing the diverse, intersectional rehabilitation goals of people living with SCI.

## Main Text

### INTRODUCTION

Empirical data and computational modeling support the view that electrical spinal stimulation nonspecifically increases net spinal sensorimotor excitability via direct recruitment of large diameter, low-threshold afferent fibers and transsynaptic activation of low-threshold interneurons (*1–9*). This diffuse excitation is sculpted into functionally relevant movement commands by inhibitory proprioceptive networks intercalated amongst spinal motor pools prior to its integration by motoneurons (*2, 5*). In this way, task specificity is not afforded by the stimulation itself, but rather by the state of the reciprocally organized propriospinal networks during stimulation-enabled movements (*2, 5*).

However, recent studies have hinted that the neuromodulatory profile of electrical spinal stimulation may be considerably more nuanced than this canonical understanding. In neurologically intact rats, direct electrical stimulation in the vicinity of spinal motor pools has revealed a different set of actions: a predictable increase in net spinal motor excitability coupled with a net decrease in high-threshold nociceptive transmission and minimal effects on low-threshold sensory transmission (*10*). Such a profile would be highly advantageous for spinal cord injury (SCI)-related applications, in which reduced voluntary motor output is frequently accompanied by pathologically elevated spinal responses to sensory feedback resulting in SCI-related neuropathic pain (SCI-NP), spasms, and spasticity (*11–14*). By differentially rebalancing spinal sensory and motor excitability to restore appropriate patterns of neural transmission in networks below the lesion, such a neuromodulatory profile would hold considerable promise for delivering multi-modal therapeutic benefits.

The extent to which motor-targeted spinal stimulation can depress high-threshold sensory transmission while preserving low-threshold sensory transmission in the chronically injured spinal cord is unestablished. Central to this question is the recognition that sensory-dominant regions of the spinal cord undergo a phenomenologically well-documented yet mechanistically enigmatic sequelae of maladaptive plastic changes during the chronification process that results in an overt state of sensory hyperexcitability (*15–17*). Indeed, the spinal ecosystem is so dramatically altered when spontaneous reinforcement of maladaptive plasticity is allowed to proceed unchecked that it remains an open question whether nociceptive specific (NS) and wide dynamic range (WDR) neurons even retain the capacity to be meaningfully downconditioned after SCI.

Here, we directly test the hypothesis that motor-targeted spinal stimulation – specifically, plasticity-promoting intraspinal microstimulation (ISMS) – depresses high-threshold nociceptive neural transmission while preserving low-threshold sensory transmission in rats with moderate to severe sensorimotor impairments secondary to chronic SCI. Using microelectrode arrays spanning the dorso-ventral extent of gray matter of the lumbar enlargement, we access large populations of discrete and functionally diverse neurons *in vivo.* This approach enables us to quantify the discharge characteristics of individual NS and WDR neurons while simultaneously inducing natural sensory feedback from the periphery before, during, and after delivery of motor-targeted ISMS. These studies provide a translational roadmap for developing a new class of spinal stimulation-based therapies specifically intended to afford multimodal therapeutic benefits – here, applied (but not limited) to simultaneous rehabilitation of movement impairments and SCI-NP.

## RESULTS

We characterized the potential effects of therapeutic motor-targeted ISMS on spinal sensory transmission *in vivo* in 15 rats with moderate to severe hindlimb paralysis at least 6 weeks following a midline T8/T9 spinal contusion injury. Seven of the animals exhibited nociceptive withdrawal reflexes and cognitive responses consistent with SCI-NP, 7 of the animals did not exhibit behavioral signs of SCI-NP, and 1 animal lacked sufficient ability to withdraw the hindlimb from a stimulus and therefore was not assessed for SCI-NP. Microelectrode arrays (MEA) were used to access the dorso-ventral extent of the L5 spinal segment (**Fig. 1A**), which enabled extracellular neural activity to be recorded simultaneously from sensory-dominant, motor-dominant, and sensorimotor integrative regions of the gray matter. This approach also permitted acquisition of neural activity during concurrent delivery of ISMS as well as during periods of natural nociceptive and non-nociceptive sensory feedback. ISMS was delivered as open-loop single pulses of charged balanced current at 7Hz and 90% of resting motor threshold to the electrode with the lowest threshold for evoking a twitch. Sensory feedback was induced by application of graded pressure to the most sensitive receptive field of the glabrous skin of the hindpaw ipsilateral to the implanted microelectrode array.

**Figure 1.**
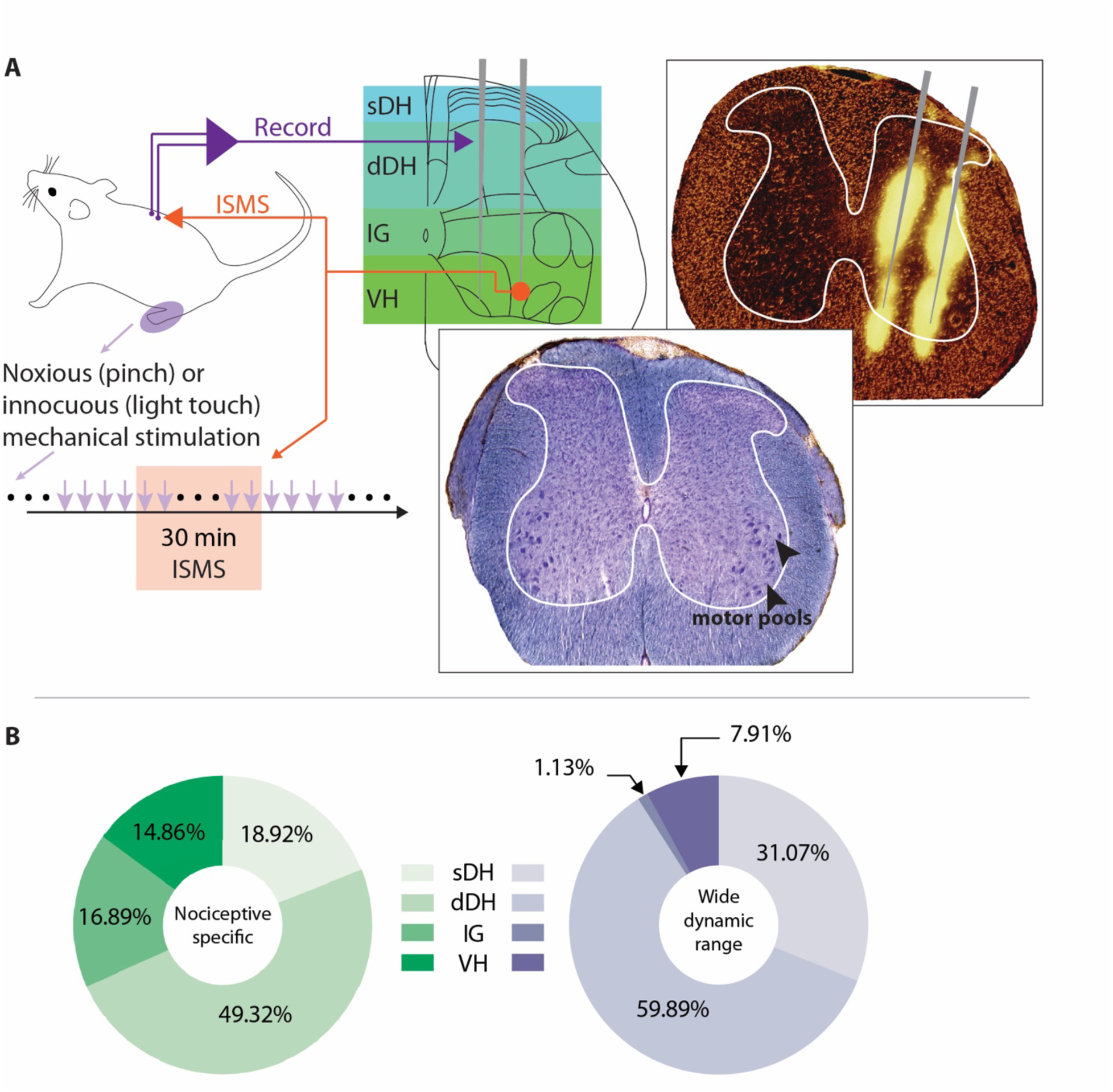
Intraspinal microelectrode arrays enable *in vivo* characterization of sensorimotor neural transmission during motor-targeted spinal stimulation. (**A**) Clockwise from top: 32-channel microelectrode arrays (MEA) were implanted perpendicular to the midline at the L5 dorsal root entry zone. MEA coverage extended from the sensory-dominant dorsal horn to the motor-dominant ventral horn; superficial dorsal horn (sDH), deep dorsal horn (dDH), intermediate gray (IG), and ventral horn (VH). Top right: histological image of electrode tracks, with fluorescent-labeled MEA illuminated by RFP filtered light (EVOS RFP 2.0 Light Cube, wavelength: Ex: 542/20 Em: 593/40; ThermoFisher Inc.,). Bottom center: histological image double-stained for cell bodies and myelin, highlighting the location of motor pools. Bottom left: mechanical stimulation was delivered to the plantar surface of the ipsilateral hindpaw before, during and after sub-motor threshold ISMS. (**B**) Anatomical distribution of functionally classified neurons (*N* = 15 rats). Green: nociceptive specific neurons; purple: wide dynamic range neurons.

Spinal responses to sensory feedback before, during, and after ISMS were based on the discharge characteristics of individual phenomenologically identified neurons. Specifically, a wavelet-based spike sorting algorithm (*18*) decomposed multi-unit extracellular neural activity into a series of spike times of discrete neurons. Those neurons were then classified as wide dynamic range (WDR) or nociceptive specific (NS) based upon their sensitivity to nociceptive and non-nociceptive sensory transmission. Neurons exhibiting increased discharge rates during both modalities of neural transmission were classified as WDR. Neurons whose discharge rate increased only during nociceptive transmission were classified as NS. All other neurons were termed non-classified (NC), regardless of whether they were tonically or phasically active.

On average, we identified 10 (±2) NS neurons and 12 (±1) WDR neurons per animal. We also identified an additional 50 (± 4) well-isolated yet functionally non-classified (NC) neurons per animal. These neurons were distributed throughout the dorso-ventral extent of the L5 gray matter. We broadly categorized their locations as falling into one of four regions: superficial dorsal horn (sDH, located ∼100-300 μm deep to the dorsal surface of the spinal cord); deep dorsal horn (dDH, located ∼400-1000 μm deep); intermediate gray (IG, located ∼1100-1300 μm deep); and ventral horn (VH, located ∼1400-1600+ μm deep). The average anatomical distribution of NS neurons per animal was sDH: 2 (±1), dDH: 5 (±1), IntZ: 2 (±1), and VH: 1 (±1); for WDR neurons, the average anatomical distribution was sDH: 4 (±1), dDH: 7 (±1), IntZ: 0, VH: 1 (±1); and for NC neurons the distribution was sDH: 3 (±1), dDH: 19 (±2), IntZ: 15 (±1), VH: 13 (±1) neurons (**Fig. 1B**).

### Motor-targeted ISMS depresses nociceptive transmission in the injured spinal cord

In neurologically intact rats, brief periods of motor-targeted ISMS – that is, ISMS delivered directly in the vicinity of spinal motor pools – can immediately and persistently modulate intraspinal nociceptive transmission (*10*). However, it is difficult to generalize these findings to sensorimotor networks below an SCI given the maladaptive plasticity and profound alterations in neural excitability that occur during the chronification process. Such excitability changes are illustrated in **Fig. 2A**, which shows spinal responses to nociceptive and non-nociceptive sensory transmission in a representative rat without SCI (left panel), a rat with chronic SCI (but no behavioral signs of SCI-NP; middle panel), and a rat with chronic SCI-NP (right panel). Increased discharge rates are clearly visible in the two animals with chronic SCI, as indicated by a shift towards warmer tones in the color spectrum; further exacerbations are evident in the animal with SCI-NP. Additionally, the spatial distribution of spinal responses to sensory feedback increases dramatically in the animals with chronic SCI, consistent with an overall state of dorsal horn hyperexcitability secondary to disinhibition due to a lack of descending neuromodulatory drive.

**Figure 2.**
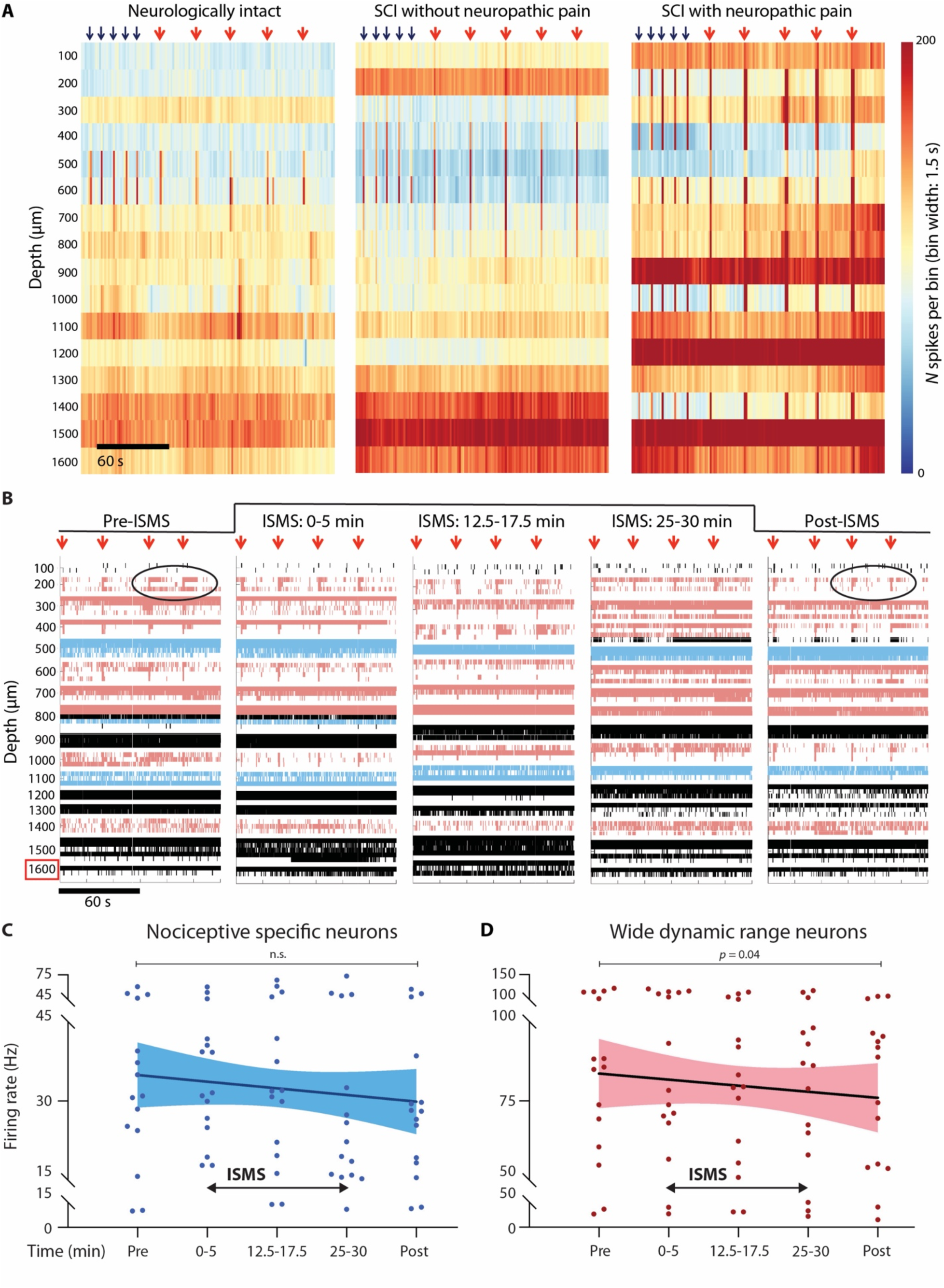
Motor-targeted ISMS leads to widespread modulatory actions that counter sensory hyperexcitability post-SCI. (**A**) Time-resolved histograms of multi-unit spiking activity across one shank of the microelectrode array in 3 representative rats during natural sensory transmission. Warmer colors indicate more spikes per unit time; cooler colors indicate fewer spikes. Black arrows above the plots indicate times of innocuous light touch of the glabrous skin of the ipsilateral hindpaw; red arrows indicate times of noxious pinch of the same receptive field. Periods between arrows depict spontaneous neural transmission (i.e., no induced sensory transmission or ISMS). Note the clear increase in the amount and anatomical spread of spiking activity during induced sensory transmission in the animals with SCI (and in particular SCI-NP) relative to the neurologically intact animal. This phenomenon is a hallmark of sensory hyperexcitability post-SCI. (**B**) Raster plot from a single rat depicting spiking activity of individual well-isolated neurons before, during, and after motor-targeted ISMS. Red arrows above the subplots indicate times of noxious pinch of the glabrous skin of the ipsilateral hindpaw. Neurons shaded in blue were determined to be nociceptive specific (NS) and neurons shaded pink were wide dynamic range (WDR). Motor-targeted ISMS was delivered on the bottom-most electrode (highlighted via red box) during the times indicated above the raster plots. Black circles highlight the reduction of sustained discharge in response to instances of noxious peripheral stimuli that was often evident following ISMS. (**C,D**) Animal-level pooled peak discharge rates for all NS (C) and WDR (D) neurons during noxious pinch of the ipsilateral hindpaw relative to ISMS duration (*N* = 15 rats). Linear regression lines of best fit are shown with accompanying 95% confidence bounds. One-way repeated measures ANOVAs with Bonferroni multiple comparisons correction were used to investigate potential differences in discharge rate across timepoint.

Thus, we first asked whether an overall cohort-level neuromodulatory effect was present across the populations of NS and WDR neurons (respectively). **Figure 2B** depicts a raster plot representation of one electrode shank before, during, and following a single 30 min trial of motor-targeted ISMS in an animal with chronic SCI-NP (the same animal as Fig. 2A, right panel). WDR neurons and NS neurons are indicated in red and blue, respectively. Pooling analogous data across all trials and animals indicated that the overall mean discharge rate of NS neurons during periods of induced nociceptive transmission was statistically invariant to ISMS (*p* = 0.22; **Fig. 2C**, left). In contrast, WDR neurons exhibited a statistically significant overall reduction in discharge rate during ISMS that remained depressed after cessation of stimulation (*p* = 0.04; **Fig. 2C**, right). This trend in WDR neurons was qualitatively similar to that previously reported in neurologically intact animals (*10*).

Previous work has shown that population-level neural trends (or lack thereof) often belie easily separable subgroups that reveal more granular insights into the state of spinal networks (*10, 19–23*). Thus, we speculated that for both the NS and the WDR populations, it would be possible to identify subpopulations that exhibited either ISMS-induced potentiation or depression of discharge rate during nociceptive transmission. However, it was also reasonable to question whether the elevated state of dorsal horn excitability common to chronic SCI (e.g. **Fig, 2A**) would constrain the modulatory capacity of each population. Such an effect could lead to a lack of identifiable subgroups, whether because discharge rates were indeed invariant to ISMS or because all neurons within a population converged towards a similar response to stimulation.

To address this question, we attempted to divide the NS and WDR neurons into subpopulations that were either potentiated by ISMS (post-ISMS discharge rate > pre-ISMS discharge rate) or depressed by ISMS (post-ISMS discharge rate < pre-ISMS discharge rate) (**Figs. 3** and **4** for NS and WDR neurons, respectively). Assignment of a neuron to the ‘potentiated’ or ‘depressed’ category was based solely on that neuron’s post-ISMS mean discharge rate relative to its pre-ISMS mean discharge rate during nociceptive transmission; changes in discharge rate *during* ISMS did not influence the classification. Thus, it was possible that a neuron coded as potentiated based on its pre/post discharge rates could exhibit *decreased* discharge rates during ISMS. Likewise, a neuron coded as depressed could have exhibited increased discharge rates during ISMS, even though its post-ISMS rates were lower than its pre-ISMS rates.

**Figure 3.**
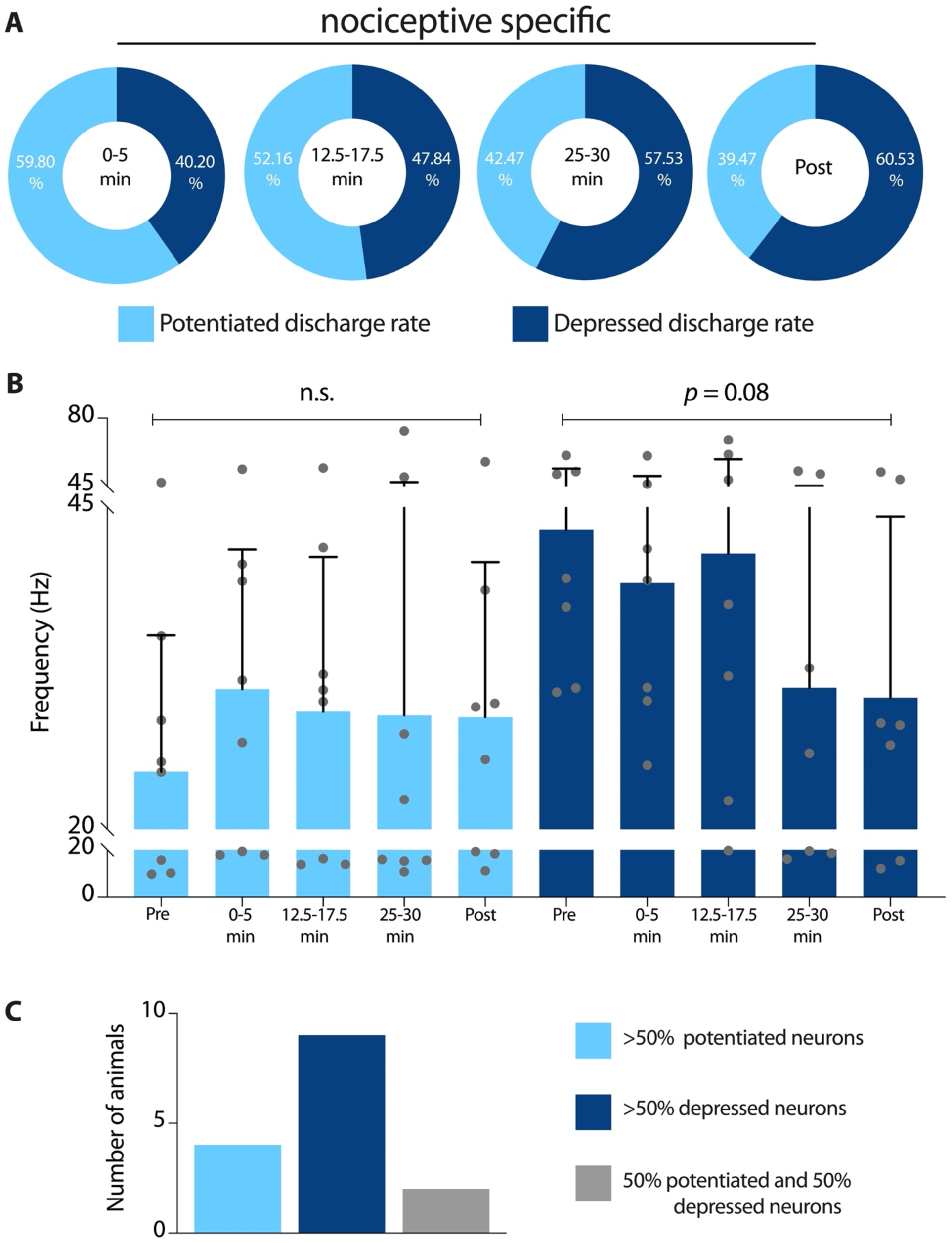
Motor-targeted ISMS drives progressive decreases in the responsiveness of nociceptive specific neurons to noxious peripheral stimuli. (**A**) Proportion of identified NS neurons exhibiting potentiated (light) or depressed (dark) firing rates during induced nociceptive transmission relative to duration of motor-targeted (*N*=15 rats). (**B**) Mean discharge rate (± sem) of potentiated (light) and depressed (dark) NS neurons, respectively, during induced nociceptive transmission relative to ISMS. Potential changes in discharge rate across time were assessed via one-way repeated measures ANOVA with Bonferroni correction for multiple comparisons. (**C**) Number of animals in which the majority of NS responses were potentiated (light), depressed (dark), or evenly divided (gray) during nociceptive transmission after ISMS relative to before ISMS.

**Figure 4.**
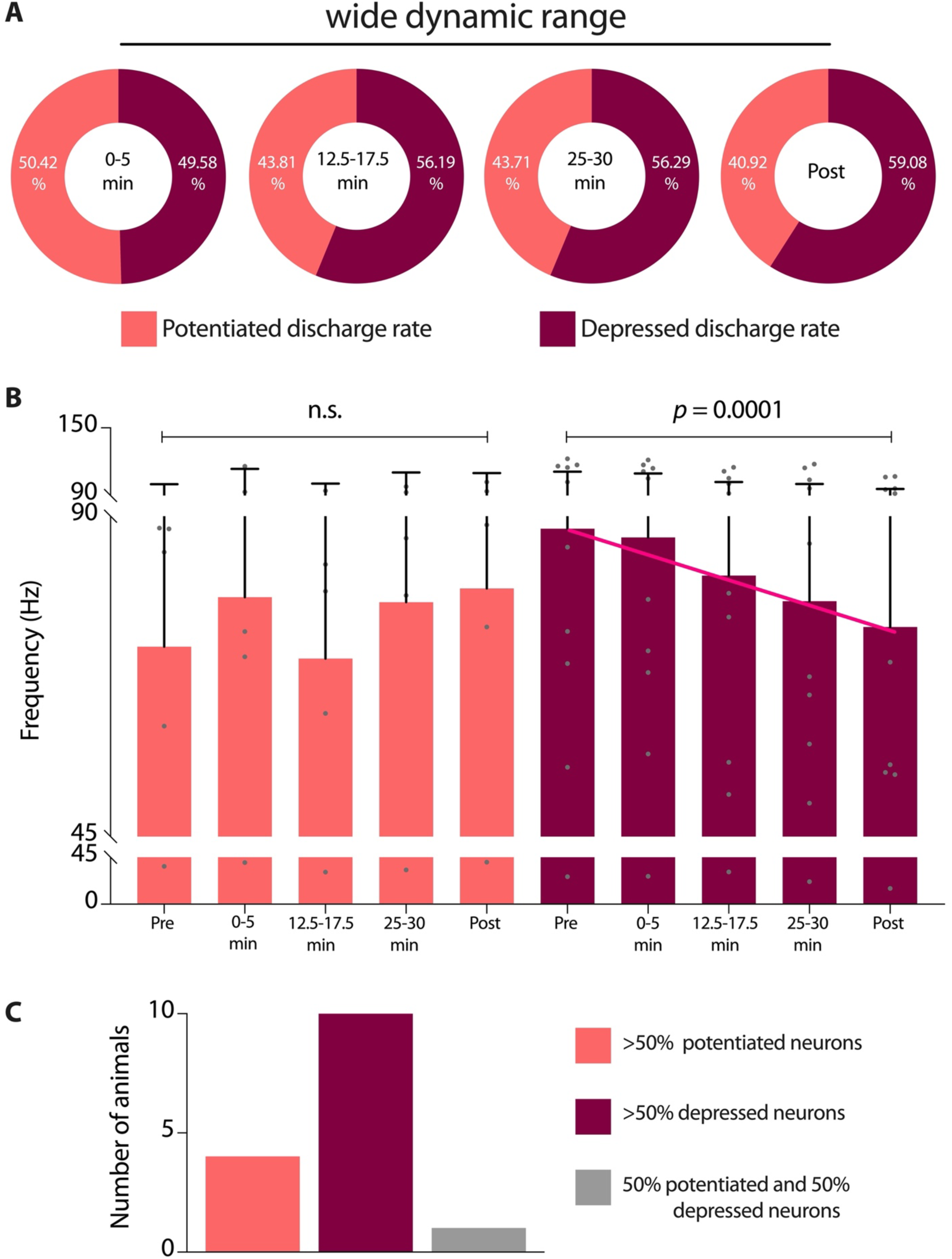
Motor-targeted ISMS drives progressive decreases in the responsiveness of wide dynamic range neurons to noxious peripheral stimuli. (**A**) Proportion of identified WDR neurons exhibiting potentiated (light) or depressed (dark) firing rates during induced nociceptive transmission relative to duration of motor-targeted (*N*=15 rats). (**B**) Mean discharge rate (± sem) of potentiated (light) and depressed (dark) WDR neurons, respectively, during induced nociceptive transmission relative to ISMS. Potential changes in discharge rate across time were assessed via one-way repeated measures ANOVA with Bonferroni correction for multiple comparisons. (**C**) Number of animals in which the majority of WDR responses were potentiated (light), depressed (dark), or evenly divided (gray) during nociceptive transmission after ISMS relative to before ISMS.

This approach was chosen to avoid biasing the entire population-level ‘during-ISMS’ statistics. If neurons were reclassified at each timepoint during ISMS, some neurons would oscillate between the potentiated and depressed groups as ISMS continued. This would force a divergence between the two subpopulations that could inflate the statistical analyses. However, as a result of this choice, our estimates of the proportion of neurons with decreased discharge rates during ISMS is likely an underestimate of the instantaneous dynamics of the population. The same holds true for the mean discharge rate of the depressed subpopulations at the corresponding timepoints. Thus, our choice results in a conservative estimate of discharge rate depression.

Across animals, the subpopulation of NS neurons exhibiting depressed responses to nociceptive transmission increased with increasing ISMS duration, obtaining a majority prior to discontinuation of stimulation (**Fig. 3A**). In tandem, the average discharge rate of these neurons progressively decreased, albeit somewhat modestly (**Fig. 3B**, *p* = 0.08). Notably, however, the proportion of NS neurons exhibiting depressed firing rates during ISMS persisted effectively unchanged following discontinuation of ISMS. This outcome was unexpected, as the proportion of ISMS-depressed NS neurons in a recently reported cohort of neurologically intact rats exhibited a ∼25% decrease upon cessation of 30 min of stimulation. More surprisingly still, the proportion of NS neurons exhibiting persistent depression after discontinuation of ISMS was also greater in animals with chronic SCI (∼61%) than that reported in animals without neurological injury (∼55%) (*10*). Together, these findings reveal unexpectedly robust carryover effects.

Regarding the subpopulation of NS neurons potentiated by ISMS, three features of bear noting. First, the average peak discharge rate of these neurons was realized within the first 5 min of ISMS and remained unequivocally stable for the duration of stimulation (**Fig. 3B**, *p* = 0.27). This finding implies that ISMS did not contribute to unintended windup of nociceptive transmission through NS neurons, which could have exacerbated the behavioral consequences of SCI-NP. Second, the number of animals in which the majority of NS neurons was depressed by ISMS was more than double the number of animals in which the majority were potentiated (9 vs. 4; **Fig. 3C**). And third, the average peak discharge rate of potentiated NS neurons was lower than that of the complementary subpopulation depressed by ISMS. It is not clear whether this finding reflects a distinction between the cell types in the two phenomenologically classified groups (i.e., the potentiated vs. depressed NS neurons) or if it is merely coincidental. Together, these findings reinforce the conclusion that ISMS did not drive an overall increase in net nociceptive transmission.

WDR neurons were evenly divided between potentiated and depressed subpopulations during the first 5 min of ISMS (**Fig. 4A**). Unlike NS neurons, however, the highest proportion of ISMS-depressed WDR neurons (∼56%) was realized within the first 10-15 min of stimulation and remained constant thereafter. This finding suggests that the depressive effect of ISMS on WDR responses did not habituate over the course of stimulation, a conclusion supported by the observation that peak discharge rate progressively decreased throughout the duration of ISMS (**Fig. 4B**, *p* = 0.0001). The proportion of WDR neurons depressed by ISMS also remained effectively unchanged following discontinuation of stimulation, presumably reflective of depressed synaptic transmission, changes in intrinsic excitability of WDR neurons, and/or an altered mixture of endogenous neuromodulation (*10*). And although the overall proportion of WDR neurons remaining depressed after ISMS (∼60%) was less than the ∼73% previously reported in neurologically intact animals (*10*), the number of animals exhibiting net depression of WDR responses to nociceptive transmission was more than 2-fold higher than animals exhibiting a net potentiation (10 vs. 4; **Fig. 4C**). Considering also the net depressive effect of ISMS on NS neurons, these findings establish a heretofore unreported multimodal neuromodulatory effect of motor-targeted spinal stimulation: namely, concurrent depression of spinal nociceptive transmission.

### Sub-motor threshold ISMS does not robustly modify spinal responses to non-nociceptive sensory feedback, revealing unexpected modal specificity

Previous studies have hypothesized that recruitment of non-nociceptive sensory afferent pathways is necessary (albeit not sufficient) for electrical spinal stimulation paradigms to enhance voluntary motor output and, separately, to reduce nociceptive transmission. In the context of motor control, recruitment of non-nociceptive sensory afferent pathways is presumed to augment or replace weakened or absent excitatory synaptic drive to motoneuron pools below the SCI (*2, 5, 7, 24*). In the context of pain management, activation of low-threshold afferent pathways has been theorized to engage a gating-like mechanism that meters the transmission of nociceptive information through WDR neurons (*4, 8*). Yet, it has also been reported in the uninjured spinal cord that sub-motor threshold ISMS delivered amongst spinal motor pools exerts a negligible impact on WDR responsiveness to non-nociceptive cutaneous and/or proprioceptive sensory feedback (*10*).

It is unclear whether motor-targeted ISMS alters the responsiveness of WDR neurons to non-nociceptive cutaneous feedback below a chronic SCI. Therefore, we characterized the discharge rate of WDR neurons in response to such feedback before and after 30 minutes of motor-targeted ISMS (in 159 discrete WDR neurons across 14 animals). Across the entire population of identified WDR neurons, we found no difference in discharge rate following ISMS compared to the corresponding pre-ISMS baseline (pre-ISMS: 73.55 ± 4.83 Hz; post-ISMS: 76.04 ± 6.02 Hz). Consistent with this observation, the proportions of WDR neurons exhibiting potentiated and depressed responses (respectively) to non-nociceptive sensory feedback following ISMS were indistinguishable (potentiated: 49.48%; depressed: 50.52%; **Fig. 5A**).

**Figure 5.**
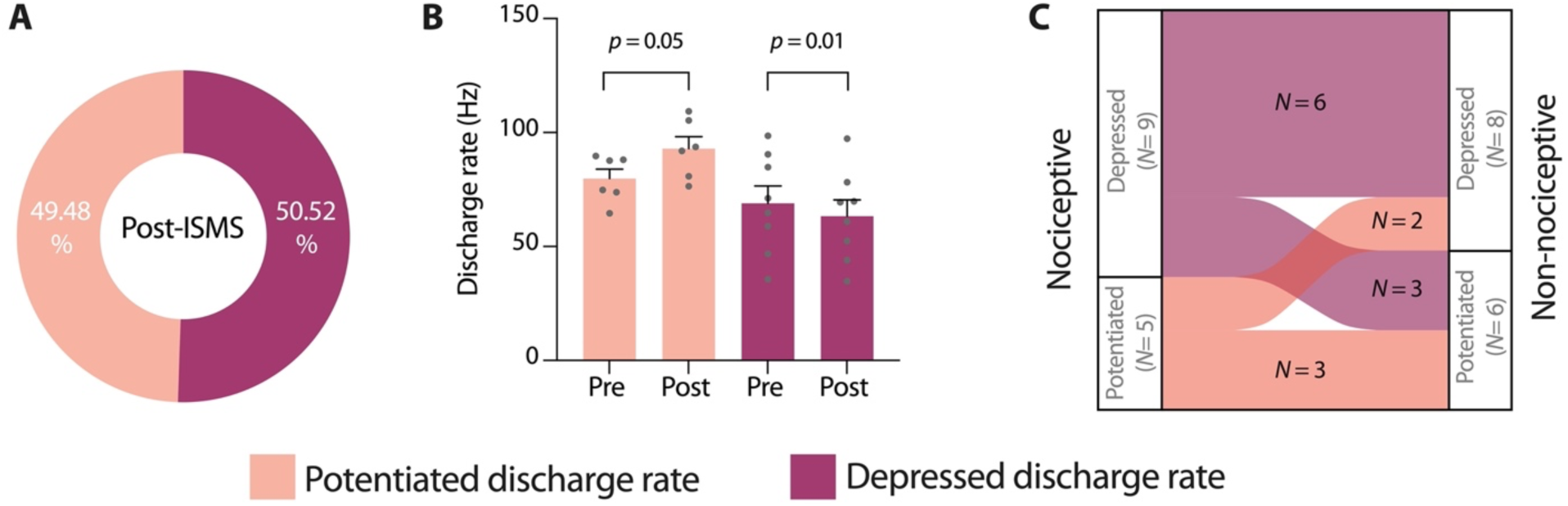
Motor-targeted ISMS exhibits an unexpectedly diminished capacity to modulate non-nociceptive cutaneous transmission relative to nociceptive transmission. (**A**) Percentage of WDR neurons (of 159 neurons across 14 rats) with potentiated (lighter) or depressed (darker) discharge rates during innocuous touches of the ipsilateral hindpaw after 30 min of ISMS relative to before ISMS. (**B**) Mean discharge rate (± sem) of WDR neurons per animal grouped by those with a majority of neurons potentiated (light) or depressed (darker color) during innocuous touches of the ipsilateral hindpaw after 30 min of ISMS relative to before ISMS. *P*-values derived from paired, 1-tailed t-tests. (**C**) Alluvial plot depicting cases in which the modulatory effects of ISMS shifted from potentiation to depression (or vice versa) when transitioning between sensory modes.

These findings suggested that motor-targeted ISMS exerted modality-specific modulatory actions that preferentially influenced nociceptive transmission relative to non-nociceptive transmission (as opposed to non-specifically modifying overall WDR excitability). But to determine if the modulatory actions of ISMS on multi-modal sensory transmission were more nuanced than they initially appeared, we attempted to subdivide the neuron-level population into animal-level groups in which the majority of WDR neurons identified in a given animal was either potentiated or depressed by ISMS. Unsurprisingly given this particular subgroup analysis, the mean discharge rates after ISMS were statistically different than those before ISMS for each of the two subpopulations (*p*=0.05 for potentiated; *p*=0.01 for depressed; **Fig. 5B**). However, in keeping with the even distribution of potentiated vs. depressed cells, the magnitude of discharge rate changes for each group was modest, averaging ∼12% when pooled across the two subpopulations.

We then sought a different approach to understand the apparent modal specificity of ISMS. We reasoned that modal specificity could manifest in either of two ways: (1) differential modulatory actions across modes of sensory transmission (e.g., one mode potentiated, one mode depressed), or (2) an enhanced modulatory capacity during one mode of sensory transmission relative to another. To this end, we first determined the number of animals in which the net modulatory action of ISMS shifted from potentiation to depression (or vice versa) when transitioning between nociceptive and non-nociceptive transmission. In total, 5 (of 14) animals exhibited such behavior, with three animals having depressed responses to nociceptive transmission and potentiated responses to non-nociceptive transmission, and 2 animals manifesting the opposite trend (**Fig. 5C**).

Next, we quantified the average magnitude of discharge rate changes associated with ISMS during nociceptive and (separately) non-nociceptive transmission in each animal. Across all animals, and regardless of whether a given modality was depressed or potentiated, we found a 3-fold greater change in discharge rate during nociceptive transmission compared to non-nociceptive transmission. In only 1 animal was the magnitude of change in non-nociceptive transmission demonstrably greater than during nociceptive transmission, and in 2 animals the change in ISMS-associated discharge rate was indistinguishable across modes (both cases in which non-nociceptive transmission was potentiated on balance).

These findings could not be explained by a ‘floor’ effect, wherein discharge rates during non-nociceptive transmission prior to ISMS were low enough that they could not be reduced further. Indeed, the mean discharge rates across animals for WDR responses to nociceptive and non-nociceptive transmission were indistinguishable prior to ISMS (*p* = 0.24; 2-tailed t-test). Thus, in 11 of 14 animals the modulatory actions of ISMS were less robust during non-nociceptive transmission than during nociceptive transmission, leading to the conclusion that the apparent modal specificity of ISMS is predominantly the result of preferential modulation of nociceptive transmission and not differential modulation of each mode.

### Overall dorsal horn excitability is not modulated by motor-targeted ISMS despite the modulatory actions on nociceptive transmission

Finally, it was reasonable to envision a scenario in which broad shifts in overall spinal or regional neural excitability associated with ISMS led to differential modulation of nociceptive vs. non-nociceptive responsiveness. Such a finding would argue against the explanation that ISMS intrinsically (albeit enigmatically) drives modality specific effects and would instead suggest that this observation was an epiphenomenon related to a complex, emergent pattern of network activity. In either case, however, it is worth noting that the effect itself – net depression of nociceptive transmission coupled with negligible impacts on non-nociceptive sensory transmission – would be highly advantageous in the context of neuromodulatory therapies for people living with SCI, particularly SCI-NP and/or spasms.

To determine if diffuse, pseudorandom changes in neural excitability were associated with ISMS, we considered the population discharge rate of *all* identified neurons from each animal – NS, WDR, and non-classified – during periods of spontaneous neural transmission before and after ISMS. We began by investigating the composite discharge rate spanning the dorso-ventral extent of the L5 spinal segment (i.e., including the SDH, DDH, IG, and VH) to gauge neural excitability at the broadest level. After a modest but statistically significant segment-wide increase in discharge rate during the first 5 min of ISMS (pre-ISMS: 6.11 ± 0.48 Hz; first 5 minutes of ISMS: 7.12 ± 0.45; *p* = 0.003; **Fig. 6**), the net increase in excitability habituated and became indistinguishable from pre-ISMS baseline levels for the remainder of stimulation and following cessation of ISMS.

**Figure 6.**
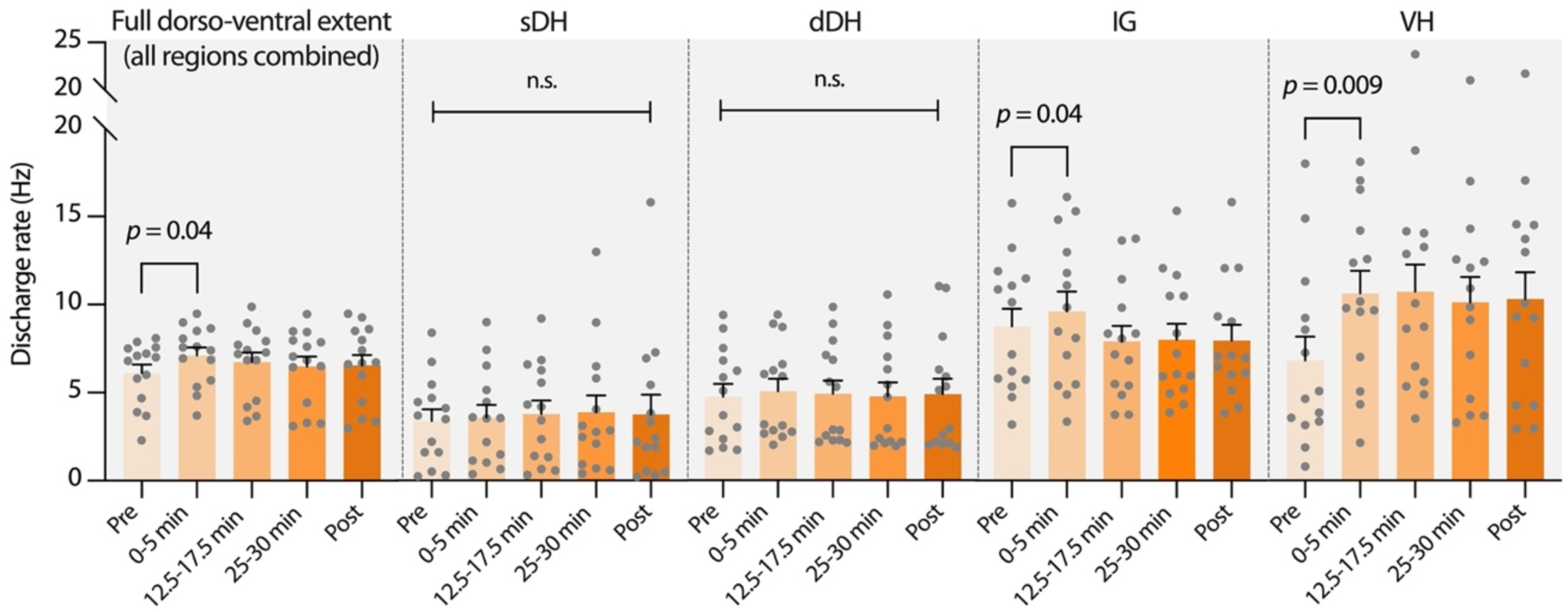
Modulatory actions of ISMS on nociceptive transmission are not accompanied by broad shifts in segment-wide neural excitability. Bar plots depict the mean discharge rate of all active neurons (NS, WDR, and NC) per animal before, during, and after motor-targeted ISMS. ISMS was delivered within the ventral horn for 30 min at sub-motor threshold intensity. No changes in discharge rate are evident in the sensory-dominant sDH or dDH, and the IG, a sensorimotor integrative region, experiences only a transient increase in excitability upon stimulation onset. By comparison, ventral horn discharge rate is increased more robustly as expected for motor-targeted ISMS. Error bars depict mean + sem; *N* = 14 animals; one-way repeated measures ANOVA with Bonferroni correction.

Next, we explored the potential impact of ISMS on region-specific excitability (i.e., considering the sDH, dDH, IG, and VH separately). We found no changes in mean spontaneous population discharge rate before or after ISMS in either the sDH or the dDH (**Fig. 6**), regions predominantly associated with transmission of nociceptive and non-nociceptive cutaneous sensory feedback. In contrast, significant differences in population discharge rate were evident within the first 5 minutes of ISMS in the IG (pre-ISMS: 8.76 ± 1.00 Hz; first 5 minutes of ISMS: 9.63 ± 1.12 Hz, *p* = 0.04) and the VH (pre-ISMS: 6.82 ± 1.35 Hz; first 5 minutes of ISMS: 10.60 ± 1.30 Hz, *p* = 0.009) (**Fig. 6**). In the IG, population discharge rate rapidly returned to its pre-ISMS baseline. In the VH, by comparison, mean population discharge rate remained elevated for the duration of ISMS and afterwards, but did not achieve statistical significance due to an accompanying increase in dispersion of the discharge rates across animals. The increased discharge rate in the VH is expected given that ISMS was delivered in this region and would be consistent with enhanced voluntary motor output as reported previously using these parameters (*25*). Ultimately, the results of these analyses indicated that it is highly unlikely that a net shift in spinal excitability contributed to the preferential depression of nociceptive rather than innocuous sensory transmission.

## DISCUSSION

Here, we demonstrated in a well-validated pre-clinical model of SCI that results in severe bilateral motor impairments and SCI-related neuropathic pain, that the responsiveness of neurons integral to the development and persistence of SCI-related neuropathic pain can be reduced by motor-targeted ISMS. We also show that these depressive actions are specific to nociceptive transmission, as spinal responsiveness to non-nociceptive transmission is largely unchanged by stimulation and motor-related transmission is potentiated.

It was not clear *a priori* whether sub-motor threshold ISMS parameterized to enhance voluntary motor output would be capable of altering neural transmission in sensory networks already made pathologically overactive by SCI (e.g., as in **Fig. 2A** and (*23*)). In point of fact, it was to some extent unclear whether the implicated networks would even retain the capacity for demonstrable plasticity, regardless of the specific approach used to affect such a change. After all, maladaptive neural activity was allowed to develop unchecked and to chronify naturally, resulting in a ‘worst case scenario’ for plasticity-promoting interventions. And whereas pre-clinical and clinical studies have repeatedly demonstrated the ability of motor-dominant circuits below a lesion to beneficially and durably reorganize (*25–32*), neural transmission in spinal pain pathways has been comparatively difficult to shape. Indeed, interventions explicitly designed to promote functionally-relevant neural plasticity as well as those that attempt to pharmacologically rebalance spinal excitability have been met with variable and generally nominal success – especially when initiated after establishment of the neuropathic pain state (*33, 34*).

Thus, it was reasonable to speculate that reinforcement of inappropriate patterns of neural transmission during the chronification process, combined with a lack of descending inhibition, would have rendered the spinal environment impermissive for allowing focal, low-amplitude stimulation delivered distantly amongst the motor pools to modulate transmission in spinal pain pathways. Alternatively, it was also reasonable to predict that injection of *any* current into the spinal cord below the lesion would increase segment-wide excitability, exacerbating overactive spinal responses to sensory feedback by trans-synaptically depolarizing many of the same neurons (or their inputs) responsible for *de novo* development of the hyperexcitable state. Yet, despite this potent backdrop of excitability, ISMS rapidly modulated transmission in spinal pain pathways, resulting in depressed NS and WDR responses to nociceptive transmission. Although the present stimulation parameters (frequency, location, duration, etc.) were not designed to promote enduring plasticity nor was the study designed to systematically characterize the washout timecourse of carryover effects, that we observed persistent excitability changes at all – and with only a single, relatively short session of ISMS – indicates both that the capacity for plasticity remains intact and that it has the potential to be unlocked via spinal stimulation.

A particularly intriguing aspect of these findings was the differential impact of ISMS on spinal networks predominantly subserving different functions. Potentiated transmission in motor-dominant ventral horn networks was evident as expected (e.g., **Fig. 7**), but this excitation was coupled with net depression of nociceptive transmission and effectively unchanged transmission of non-nociceptive cutaneous sensory feedback. More interestingly, these actions were defined across functional but not anatomical boundaries. Indeed, the overall invariance of spinal segment-level excitability during ISMS could not be explained by opposing excitatory and inhibitory actions across anatomical regions, effectively cancelling the appearance of a segment-wide effect. For example, depressed nociceptive transmission was evident in NS and WDR neurons identified in the IG and VH, regions that experienced a net increase in excitability during ISMS, whereas the responsiveness of NS and WDR neurons in the sDH and dDH was also depressed by ISMS, yet these anatomical regions saw no net change in excitability.

The mechanistic underpinning of these findings is unknown, and caution should be exercised in their interpretation. Indeed, the number of functionally identified NS and WDR neurons was substantially lower than the total number of well-isolated neurons identified in each animal (which included functionally non-classified neurons). It is also possible that the balance of excitation and inhibition would shift under awake, behaving conditions. For example, it has been elegantly demonstrated that appropriate recruitment of pre-motor proprioceptive networks is essential for providing the inhibition necessary to sculpt the diffuse excitation mediated by epidural spinal stimulation (*2, 5*). And although it is incompletely understood whether analogous engagement of supporting circuits could play a role in shaping the balance of excitation and inhibition within sensory-dominant networks, it would seem a plausible, if not likely, scenario; many examples of natural sensory gating phenomena have been reported (*35–38*). In fact, it is possible that such a mechanism could have contributed to what phenomenologically appeared to be heterosynaptic plasticity in WDR neurons, wherein long-term depression of high-threshold inputs was contrasted by negligible changes at low-threshold synapses converging onto the same cells. Regardless, if borne out by future work in awake, behaving animals, the modulatory actions observed here would distinguish sub-motor threshold ISMS from epidural spinal stimulation and provide a powerful new platform for developing multi-modal rehabilitation therapies.

The role of the ISMS protocol itself should also be underscored. In addition to broadly demonstrating that the input-output function of pain-related spinal neurons can be modified in the chronically injured spinal cord, the results also specifically demonstrate that electrical spinal stimulation has the capacity to effect such a change. Two particular aspects of the stimulation protocol bear noting.

First, stimulation was delivered in an open-loop fashion rather than in an activity-dependent manner. That is, delivery of ISMS was uncorrelated with ongoing neural activity. As a result, the ISMS paradigm made no explicit attempts to restore or reinforce natural, appropriate patterns of neural transmission in the modulated networks. Previously, it has been shown that activity-dependent spinal stimulation leads to functional gains that far exceed those afforded by open-loop stimulation, presumably by more optimally engaging circuits below the injury to promote beneficial neural plasticity (*25, 27, 39, 40*).

Pre-clinical models also indicate that the enhanced efficacy of activity-dependent stimulation is not immediate. Rather, the effects of activity-dependent stimulation paradigms diverge from open-loop paradigms several weeks into an intervention, continuing to enhance function beyond open-loop approaches over time and leading to therapeutic gains that can persist for weeks after discontinuation of stimulation (*25, 40*). Together, these observations suggest that the plasticity observed here may be a conservative representation of what could be achieved with activity-dependent stimulation and/or in the context of a directed rehabilitation intervention.

Second, modulation of sensory neurons was realized with *motor-targeted* ISMS. Indeed, the approach was not configured to directly alter the input-output function of dorsal horn neurons, for example by juxtaposing the stimulating electrode in a region dense with identified NS or WDR neurons. Instead, ISMS was delivered adjacent to spinal motor pools in the ventral horn. This distinction reveals in a well validated pre-clinical model of SCI that it may be possible to purposefully engineer a spinal stimulation-based intervention to deliver multi-modal therapeutic benefits. Here, those benefits centered upon simultaneously rebalancing the pathologic sensory and motor transmission that contribute to debilitating motor impairments and SCI-NP. However, the high interconnectivity and diversity of physiological functions mediated by sensorimotor networks in the lumbar (or cervical) enlargement suggests that other combinations could also be envisioned. Paradigms intended to simultaneously enhance motor output while restoring bowel, bladder, or sexual function would address many of the issues routinely deemed most important by people living with SCI, for example (*41–44*).

Given that ISMS will require additional time to progress through the translational pipeline before potentially emerging as a viable clinical option, the logical question becomes how best to accelerate introduction of these ideas into advanced yet currently available therapies. Fortunately, many of the results obtained with ISMS support the idea that epidural spinal stimulation may also be capable of delivering multi-modal benefits. Nominally, it is worth reiterating that clinically available epidural spinal stimulators were originally developed to manage medically refractory pain conditions, and the parameter sets and electrode montages used in early motor rehabilitation applications closely mirrored those employed by the pain field. In fact, it has traditionally been posited that the analgesic effects of paresthesia-based spinal stimulation paradigms are mediated by mechanisms highly overlapping with those intended to enhance spinal motor output (although this interpretation is now understood to be incomplete) (*2, 4, 5, 7, 8, 10, 24*). Thus, the literal ability of epidural spinal stimulation to modulate both sensory and motor networks is not in question.

A striking feature of these results was the specificity with which motor-targeted ISMS modulated spinal responsiveness to nociceptive transmission despite the broad spatial extent over which the identified neurons were located – particularly given that stimuli were delivered at sub-threshold intensity for evoking muscle contractions. This diffusivity implies that focally delivered stimulation may not be essential to achieve qualitatively similar sensory and motor effects to those observed here. At present, however, the majority of epidural stimulation paradigms to enhance motor output use electrical current intensities suprathreshold for evoking contractions, whereas epidural stimulation for pain management remains subthreshold for motor unit recruitment. And unlike suprathreshold stimulation, we did not observe net increases in non-nociceptive transmission via large-diameter afferents. Thus, to the extent that stimulation location(s) and target(s) will have implications for stimulation intensity and presumably for the ensuing modulatory actions, it will be critical to reconcile how to parameterize epidural spinal stimulation for multi-modal applications. One starting point could be renewed investigation of sub-motor threshold epidural stimulation (*9, 45*), which is designed to augment volitional attempts to move by facilitating weakened yet spared pathways rather than to produce the intended movement(s) directly.

In the immediate term, two additional considerations would accelerate translation of these results. First, ongoing or planned studies of epidural spinal stimulation intended to enhance voluntary motor output after SCI could specifically incorporate assessments of pain perception and sensory acuity into their battery of outcome measures; there are many validated instruments for these purposes. And while fewer such studies are ongoing, there is likewise much to be gained from incorporation of motor assessments into studies of epidural spinal stimulation for SCI-NP. And second, enrollment of people living with sensorimotor incomplete SCI, especially those living with SCI-NP, into trials of motor-targeted epidural spinal stimulation would provide a wealth of insights. Purposeful, systematic acquisition of data pertaining to the off-target effects of epidural spinal stimulation for motor rehabilitation will move the neuromodulation field towards conceptualization and development of therapies focusing on the complex, interrelated rehabilitation goals of people living with SCI.

## MATERIALS AND METHODS

### Study design

We studied the effects of ISMS intended to enhance motor output on spinal sensory transmission in the chronically injured spinal cord. The study included 15 adult male Sprague-Dawley rats (∼400-550 g) with chronic T8/T9 SCI. All procedures described herein were approved by the Institutional Animal Care and Usage Committee of Washington University in St. Louis.

In a terminal electrophysiological experiment at least 6 weeks after SCI, we recorded extracellular neural activity throughout the dorso-ventral extent of the L5 spinal gray matter while inducing natural nociceptive or non-nociceptive feedback before, during and after 30-minute epochs of sub-motor threshold ISMS delivered to the L5 spinal motor pools.

Trials had the following general sequence: (a) 1 min of innocuous mechanical stimulation of the receptive field, delivered as a series of light touches of the most sensitive region of the receptive field; (b) 1 minute of spontaneous activity (i.e., no mechanical stimulation of the receptive field); (c) 3-5 minutes of noxious mechanical stimulation of the receptive field, delivered as a ∼1-2 s pinch of the most sensitive region of the receptive field every 30 s; (d) 30 minutes of motor-targeted ISMS combined with phasic noxious mechanical stimulation of the receptive field; (e) 3-5 minutes of noxious mechanical stimulation of the receptive field (as above, in c); (f) 1 minute of spontaneous activity; and (g) 1 min of innocuous mechanical stimulation of the receptive field (as above, in a).

This trial structure allowed assessment of potential modulatory actions of ISMS on spinal nociceptive and non-nociceptive transmission. Inclusion of nociceptive and non-nociceptive transmission also enabled classification of neurons as nociceptive specific (NS), wide dynamic range (WDR), or other (termed ‘non-classified’, or NC). The 30 min ISMS duration was selected to mirror a previously published report of motor-targeted ISMS on spinal nociceptive transmission in neurologically intact rats (*10*). In that study, 30 min was found to be more efficacious at reducing nociceptive transmission than 2, 5, or 10 min of motor-targeted ISMS. It is also more comparable to the durations of stimulation that would be utilized for clinical pain management and household or community ambulation.

Motor targeted ISMS was delivered as a series of discrete, cathode-leading pulses at 7 Hz, 200 μs/phase, 0 s inter-phase interval, and 90% of resting motor threshold (typically ∼3-12 μA/phase). These ISMS parameters are consistent both with values that induced functionally meaningful enhancements of spinal motor output below a chronic SCI (*25*) and with stimulation intensities (e.g., 80-90% of resting motor threshold) commonly used in studies of ESS parametrized to treat pain (*46, 47*).

### Statistical analyses

All statistical analyses were based upon the instantaneous discharge rate of NS, WDR, and NC neurons during nociceptive or non-nociceptive transmission before, during, and after ISMS. For inclusion in quantitative analyses, NS and WDR neurons were required to manifest a mean pre-ISMS discharge rate >6 Hz during periods of induced nociceptive transmission and a mean discharge rate >0.11 Hz during and following ISMS; NC neurons were required to manifest a nominal mean discharge rate of >0.11 Hz throughout a given trial. These criteria ensured inclusion of only neurons that could be readily tracked across all segments of a trial, preventing artificially altering population discharge rate metrics by ‘zero-padding’.

Discharge characteristics were pooled within each animal prior to group analysis to appropriately capture the heterogeneity associated with spinal contusion injuries such as that used here. Animals were not randomized *per se*, although it was not possible to know *a priori* which animals would develop SCI-NP following injury. Behavioral analyses were conducted by an experimentalist not involved in terminal electrophysiological experiments, and it was not possible to determine during a terminal electrophysiological experiment whether a given animal did or did not exhibit behavioral signs of SCI-NP.

Mean discharge rate was defined as the grand mean of the maximum instantaneous discharge rate of each neuron across all instances of mechanical probing of the dermatome for a given timepoint. For example, if a pre-ISMS baseline epoch of nociceptive transmission contained 10 pinches of the receptive field, the peak discharge rate during each pinch was averaged for each neuron of a given type, after which those mean neuron-level rates were averaged to arrive at an animal-level grand mean.

Comparisons of population discharge rates across time (relative to ISMS) utilized one-way repeated measures ANOVAs. Separate ANOVAs were computed for NS neurons, WDR neurons, and, in the case of overall excitability, pooled NS, WDR, and NC neurons. For all comparisons, the independent variable was timepoint relative to ISMS and the dependent variable was mean discharge rate of a given neuron type per animal at that timepoint. Post-hoc analyses compared mean discharge rates before ISMS (during nociceptive transmission) to each subsequent timepoint during and following ISMS, and Bonferroni correction was used to control family-wise error rate. WDR discharge rate during non-nociceptive transmission included only pre-ISMS and post-ISMS timepoints. Thus, paired, two-tailed t-tests were used to compare these rates. Descriptive statistics presented in the text and figures display mean ± s.e.m. unless otherwise noted; specific p-values are indicated if < 0.1; otherwise, comparisons are labeled ‘n.s.’. Statistical analyses were performed using GraphPad Prism software (v9).

A power analysis was performed *a priori* (G*Power 3.1) using data extracted from a recently published report of motor-targeted ISMS on nociceptive and non-nociceptive transmission in neurologically intact rats (*10*). Those data allowed determination of a presumptive ‘best-case scenario’ effect size for ISMS-associated depression of discharge rates during nociceptive transmission. Achieved effect size in that study for one-way repeated measures ANOVAs with the same independent and dependent variables as those used here was estimated to be 0.48, which is considered a medium effect in the convention of standardized effect sizes. Using that effect size along with an α of 0.05, power (1-β) of 0.9, and a moderate correlation among repeated measures (0.6; range of 0-1, where the repeated measures are the mean discharge rate per rat across timepoints), we estimated that 7 animals would be required.

However, recognizing both the heterogeneity associated with contusion models of SCI as well as the proclivity of the dorsal horn to become hyperexcitable post-SCI, we expected to realize a smaller net change in discharge rate associated with ISMS. Thus, we chose a more conservative estimate of effect size, 0.3, which corresponds to a small standardized effect. Using this effect size instead (while carrying forward the other parameters unchanged) the required sample size was determined to be 15 animals.

### Electrophysiology

At least 6 weeks after the SCI, we performed a terminal electrophysiological procedure in each animal. A single microelectrode array (MEA) was implanted into the spinal cord at the L5 dorsal root entry zone. Each array consisted of two parallel shanks, and both shanks contained 16 discrete, vertically aligned electrodes (area: 177 μm^2^; inter-electrode spacing: 100 μm; NeuroNexus Inc., A2×16). The tips of both shanks were sharpened to aid insertion. All MEAs were custom electrodeposited with activated platinum-iridium to lower impedance and increase charge capacity (impedance: 4-10 KΩ; Platinum Group Coatings, Inc.). Prior to implantation, MEAs were coated with 1,1’-Dioctadecyl-3,3,3’,3’-tetramethylindocarbocyanine perchlorate (Sigma-Aldrich, Inc.) to aid postmortem histological localization of the electrode tracks (as in Fig. 1A, top right panel).

MEAs were coupled via an Omnetics nano connector to a Nano2+Stim headstage and Grapevine neural interface processor (Ripple Neuro, Inc.). This system enabled simultaneous recording of extracellular neural activity across all 32 MEA channels while delivering current-controlled stimulation through the MEA. It also allowed real-time visualization of multiunit neural activity across all channels. A high-fidelity electrophysiology audio amplifier with customizable equalizer (A-M Systems, Inc.) enabled further online assessment of neural transmission.

For implantation into the spinal cord, the MEA-headstage assembly was mated to a custom motorized four-axis micromanipulator with submicron resolution (Siskiyou Corp.). The MEA was then oriented perpendicular and just lateral to the midline and positioned directly above the L5 dorsal root entry zone under high magnification. Subsequently, the MEA was lowered until the bottom-most electrode on each shank came into initial contact with the dorsal surface of the spinal cord and was lightly nested amidst the dorsal roots. We then probed the glabrous skin of the ipsilateral hindpaw while monitoring dorsal root potentials in real-time with visual and audio feedback. If dermatome mapping evoked clearly correlated dorsal root potentials, MEA insertion began as detailed below. However, if no dorsal root potentials were evident during mapping or if the most responsive portion of the dermatome was not located on the plantar surface of the hindpaw, the MEA was repositioned and mapping began anew.

Once the initial implant site was established, the MEA was slowly advanced into the spinal cord. Insertion was paused every ∼25-50 μm to minimize shear and planar stress on the neural tissue, which is particularly important for preserving the viability of the small superficially located neurons in laminae I-III. When the deepest electrodes of the MEA reached the most superficial border of the deep dorsal horn (∼400-500 μm deep to the surface), we again mapped the L5 dermatome while monitoring neural transmission in real-time. If clearly correlated multiunit neural activity was evident during exploration of the desired receptive field, insertion continued. Conversely, if the most sensitive receptive field had shifted from the glabrous skin of the hindpaw to another region of the L5 dermatome, the MEA was withdrawn and repositioned. When fully inserted, the ventral-most electrodes were positioned ∼1600-1800+ μm deep to the surface of the spinal cord, corresponding to the motor pools of the ventral horn, and the dorsal-most electrodes were ∼100-200 μm deep to the surface. Once insertion was complete, we again mapped the L5 dermatome. Electrode arrays were not moved after full implantation.

### Spinal cord injury model

All SCI surgical procedures were conducted using strict aseptic techniques. Anesthesia was induced using inhaled isoflurane (∼3-4% O_2_, flow rate: 1-2 L/min) and subsequently by intraperitoneal injection (i.p.) of ketamine (80 mg/Kg) and xylazine (12 mg/Kg). Boosts of ketamine/xylazine cocktail were provided as necessary (ketamine [40 mg/Kg] and xylazine [6 mg/Kg]). After the animal was deeply anesthetized, prophylactic buprenorphine SR (1mg/Kg) was injected subcutaneously to aid post-operative analgesia.

The hair on the animal’s back was shaved and the region was cleaned 3x with alternating betadine and alcohol rubs. A ∼5 cm midline skin incision was made over the vertebral column originating approximately at the cervicothoracic junction. The dorsal musculature was dissected and retracted, and a laminectomy was performed on the T8 vertebrae under magnification (Leica Microsystems, Inc.). Vertebral clamps were secured rostral and caudal to the exposure and the animal was elevated such that its abdomen was suspended slightly above the surgical platform, minimizing respiration cycle-induced movement of the spinal cord.

We then used a dedicated rodent spinal cord impactor (Infinite Horizon Impactor, IH-04000; Precision Systems and Instrumentation, LLC) to produce a dorsal midline contusion injury. Specifically, the 2.5 mm diameter impactor tip was lowered to within ∼2 mm of the exposed, intact dura mater at the T8/T9 border before delivering an impact force of 200 Kilodynes and 0 sec of dwell time. These parameters result in moderate-to-severe hindlimb paralysis in all animals and below level SCI-related neuropathic pain in 40-60% of animals, typified by mechanical allodynia and hyperalgesia. After the contusion injury, muscles were closed in layers and the skin incision was closed with suture and surgical staples. Warmed lactated Ringer’s solution (5 mL) and antibiotic (Enrofloxacin 0.5mg/Kg) were administered subcutaneously to prevent dehydration and infections, respectively. From induction of anesthesia until closure, the SCI procedure required approximately 1-1.5 hrs.

### List of Supplementary Materials

Materials and Methods

References (*48–50*)

## Acknowledgments

N/A

## Author contributions

Conceptualization: JGM

Data curation: MB, JGM

Formal analysis: MB, JGM

Methodology: MB, JGM

Investigation: MB, JLG, JGM

Software: MB, JGM

Validation: MB, JGM

Visualization: MB, JGM

Funding acquisition: JGM

Project administration: JGM

Supervision: JGM

Writing – original draft: MB, JGM

Writing – review & editing: MB, JGM

## Data and materials availability

Data and code used in the analysis are available upon reasonable request to the authors.

## SUPPLEMENTARY MATERIALS

Materials and Methods

References (*48–50*)

### Animal care and behavioral assessments

After the SCI procedure, animals were transferred to recovery housing which included a thermal pad to maintain core temperature (∼37 °C). Heart rate, blood pressure, respiration rate, and SpO2 were continuously monitored (Kent Scientific, Inc.) until arousal. Nutritional supplements, electrolyte replenishers (Bio-Serv) and water with antibiotic (Enrofloxacin 0.5ml/Kg) and sweetener were provided to the animals during their recovery. Weight loss/gain, righting reflexes, forelimb and head/neck motor control, coat quality, demeanor, hydration, and urine composition were closely monitored for at least the first 2 post-operative. Each animal’s bladder was manually expressed ≥ 2x/day for 7 days after the injury or until voiding reflexes returned spontaneously.

We assessed all animals for behavioral signs of below-level neuropathic pain at least twice per week beginning 1-week post-SCI and continuing until the terminal electrophysiological experiment (at ≥ 6 weeks post-SCI). Common signs of SCI-related neuropathic pain in this contusion model include the presence of mechanical allodynia and/or hyperalgesia in dermatomes below the lesion. Our outcome measures included nociceptive behavioral responses as well as so-called ‘cognitive’ responses to mechanical probing of the L5 dermatomes ipsilateral and contralateral to the electrode implant.

Nociceptive responses were based on the mechanical threshold (measured in grams of force) for eliciting an aversive withdrawal reflex on the plantar surface of each hindpaw using the up-down method (*48, 49*). Forces were applied using small rigid probes, similar in profile to a micropipette tip, coupled to a von Frey system instrumented with a precision loadcell (Electronic von Frey, IITC Inc.). Ten measurements were completed in each hindpaw per session, and the highest and lowest scores were discarded; the remaining thresholds were averaged. One animal was excluded from von Frey testing because it could not voluntarily move the hindpaw from the probe due to severe motor impairment.

Cognitive responses to noxious and innocuous cutaneous sensation included vocalizations during mechanical probing of the L5 dermatome as well as attendance to the involved hindpaw as evidenced by turning the head and neck towards it and/or attempting to reach it with the forelimb(s). To reduce the well-known confounds of stoicism in prey animals, increased stress associated with temporary transfer to the behavioral testing arena, and non-pain-related hyperreflexia – all of which can alter attendance to routine handling, movements, and/or noise – rats were allowed at least 20 min to acclimate to the testing environment prior to assessment.

The presence of an SCI-NP state was established by a ≥ 50% reduction in withdrawal threshold compared to an animal’s pre-SCI baseline and/or clear cognitive signs of distress during innocuous exploration of the dermatome.

### Surgical procedure for electrophysiology experiments

Anesthesia was induced with inhaled isoflurane (∼1-3% O_2_, flow rate: 1-2 L/min), which was discontinued upon uptake of urethane (1.2 g/Kg i.p.). Deep surgical plane anesthesia was maintained with boosts of urethane as needed (0.2 g/Kg i.p.). Urethane was chosen for its ability to preserve the excitability of spinal nociceptive and sensorimotor reflex pathways (*10, 19*). After the animals were deeply anesthetized, the back and hindlimbs were shaved. Then, a ∼5 cm midline incision was made over the vertebral column on the shaved region. The exposed tissues and musculature were dissected and a bilateral T13-L2 laminectomy was performed under magnification (Leica Microsystems, Inc). For the duration of the procedure, the animal’s temperature was maintained at ∼37° C using a thermal pad, and heart rate, blood pressure, respiration rate, and SpO2 were continuously monitored (Kent Scientific, Inc.). Warmed lactated Ringer’s solution (5 mL) was administered subcutaneously every 2 hours to prevent dehydration.

After the laminectomy, animals were transferred to an anti-vibration air table (Kinetic Systems, Inc.) enclosed by a Faraday cage. Vertebrae rostral and caudal to the laminectomy were clamped with locking forceps connected to a custom, multi-degree-of-freedom fixation frame. The animals’ abdomen was raised using the fixation frame to attenuate chest cavity and spinal movements during respiration cycles. The spinal meninges were then incised rostro-caudally and reflected. The exposed spinal cord was continually bathed in homeothermic ringer solution and the MEA was implanted.

### Motor threshold determination

After MEA implantation, we determined the resting motor threshold for each animal. We delivered single pulses of charge balanced current (cathode leading, 200 μs/phase, 0 s inter-phase interval) to the electrodes located in the deepest regions of the ventral horn. Current magnitude was increased in 1 μA steps until a twitch was detected in the L5 myotome (toe twitch on ipsilateral hindpaw). Then, we reduced the current intensity in 1 μA steps until the twitch was undetectable. The lowest current intensity at which a twitch was detected, across electrodes located in the ventral horn, was taken as the resting motor threshold. The specific electrode on which the threshold was established was used to deliver ISMS on all subsequent trials.

### Spike train analysis

Raw, multi-unit neural data was pre-processed offline to remove non-physiological features (e.g., electrical noise, artifacts caused by respiration cycles) (The MathWorks, Inc.). Cleaned multi-unit data were then decomposed into spike trains of individual neurons using a well validated wavelet-based spike sorting algorithm (*10, 18, 19, 50*). Spike trains were then analyzed manually to remove any errors associated with the decomposition (e.g., predominance of interspike intervals < 2ms, non-physiological shape of the action potential or inappropriate action potential duration). Neurons not passing the manual verification stage were discarded.

